# Dissection of yeast pleiotropic drug resistance regulation reveals links between cell cycle regulation and control of drug pump expression

**DOI:** 10.1101/294579

**Authors:** Jian Li, Kristen Kolberg, Ulrich Schlecht, Robert P. St.Onge, Ana Maria Aparicio, Joe Horecka, Ronald W. Davis, Maureen E. Hillenmeyer, Colin J.B. Harvey

## Abstract

Eukaryotes utilize a highly-conserved set of drug efflux transporters to confer pleiotropic drug resistance (PDR). Despite decades of effort interrogating this process, multiple aspects of the PDR process, in particular PDR regulation, remain mysterious. In order to interrogate the regulation of this critical process, we have developed a small-molecule responsive biosensor that couples PDR transcriptional induction to growth rate in *Saccharomyces cerevisiae*. We applied this system to genome-wide screens for potential PDR regulators using the homozygous diploid deletion collection. These screens identified and characterized a series of genes with significant but previously uncharacterized roles in the modulation of the yeast PDR in addition to recapitulating previously-known factors involved in PDR regulation. Furthermore, we demonstrate that disruptions of the mitotic spindle checkpoint assembly lead to elevated PDR response in response to exposure to certain compounds. These results not only establish our biosensor system as a viable tool to investigate PDR in high-throughput, but also uncovers novel control mechanisms governing PDR response and a previously uncharacterized link between this process and cell cycle regulation.

**Significance:** Pleiotropic drug resistance (PDR) is a conserved mechanism by which cells utilize membrane bound pumps to transport chemicals out of the cell. Here, we develop a growth-based biosensor system in yeast that enables high-throughput identification of factors that transcriptionally regulate PDR. Among the novel PDR regulators identified here, we show that spindle assembly checkpoint (SAC) proteins, which are important for cell cycle regulation, inhibit hyperactivation of PDR upon drug treatment. This result provides insights into PDR regulation, as well as potential targets for therapeutic intervention, particularly in chemoresistant cancers where the cell cycle regulation is often disrupted.

## Introduction

Multidrug resistance (MDR) is a highly conserved process in mammalian, fungal, and bacterial cells that is characterized by resistance to a variety of unrelated xenobiotics (1–3). Such resistance is primarily conferred through complex interactions between a network of transcriptional regulators and genes encoding downstream transmembrane ATP binding cassette (ABC) efflux pumps (4). More than a dozen proteins in this network, also termed pleiotropic drug resistance (PDR), have been identified in *Saccharomyces cerevisiae*. Among the most extensively studied of these are plasma membrane-bound pumps Pdr5, Snq2 and Yor1 (5). Pdr5, a transporter with hundreds of verified substrates, is a functional homolog of *Candida albicans* Cdr1 and mammalian P-glycoprotein (MDR1); two transporters implicated in clinical resistance to variety of drugs (3, 6). Snq2 shares structural similarity to Pdr5 and was the first ABC transporter in yeast that was implicated in drug resistance (7), while Yor1 belongs to the Cystic Fibrosis Transmembrane Conductance Regulator (CFTR) family of transporters. All are major drivers of drug resistance in yeast (8).

Hyperactivation of the PDR is the leading cause of resistance to newly developed antifungal azole and echinocandin drugs in many fungal pathogens (3), while over expression of MDR1, the human homolog of Pdr5, creates substantial challenges in chemotherapy as drugs are pumped out of cancer cells and cannot reach therapeutic concentrations (12). As a result, molecules that modulate the activity of these pumps are among the leading candidates to overcome widespread drug resistance (13, 14).

The genetic, structural, and functional similarity of *S. cerevisiae* PDR regulators and pumps to their homologs in humans and pathogenic fungi have made yeast a valuable model to study PDR regulation, as well as mutations in the pumps themselves that lead to multidrug resistance and ABC protein diseases such as cystic fibrosis (15).

Activation of the PDR transcriptional cascade is a rapid and complex process. Extensive study has revealed Pdr1 and Pdr3, two zinc-finger transcription factors with partially overlapping roles, regulate more than half of all known pumps; Pdr5, Snq2, and Yor1 included. These regulators act through binding to conserved motifs in the promoters called PDR elements (PDRE) (4). Yrr1, another transcription factor responsive to a different set of stimuli controls the expression of Snq2 and Yor1 through binding of Yrr1-response element (YRRE) (9). Additional transcription regulators, such as Yap1 (yes-associated protein 1), Yrm1 (yeast reveromycin resistance modulator), and Msn2 (Multicopy suppressor of SNF1 mutation), have each been demonstrated to play a role in cellular response to chemical, oxidative, and hypoxic stress (4). While these transcriptional regulators are critical in determining the level of PDR expression, a detailed understanding of the entire regulatory cascade that causes xenobiotic stress to induce increased pump expression has yet to be developed. In particular, a specific compound or stress signal usually activates only a certain set of transporter genes, and for transporters with identical or highly similar promoter response elements, transcriptional response can still be quite different in response to the same stimulus (5, 10). The current set of known PDR regulators does not fully account for the specificity and diversity of PDR. Many factors that act upstream of and in conjunction with known PDR transcription factors are yet to be identified and characterized.

In order to search, on a genome-wide level, for additional proteins that participate in the transcriptional cascade responsible for drug sensing and transporter activation, we developed a novel biosensor system that couples the growth rate of yeast cells to the expression of a specific PDR transporter. We then applied this system to screens of the yeast homozygous-diploid deletion collection under drug treatment and identified deletion mutants that showed either diminished or enhanced PDR activation. Using this approach, we have discovered a series of genes with significant and previously uncharacterized roles in the modulation of the yeast PDR. Enriched among these hits are genes known to be involved in multiple areas of yeast biology, including regulation of cellular process and response to external stimuli, as well as cellular signaling and phospholipid metabolism.

Among the previously uncharacterized regulators of the PDR identified here are the Mad family of proteins involved in the mitotic SAC complex, the disruption of which has been explored as an anticancer strategy (11). We demonstrate that such disruption of mitotic spindles leads to elevated PDR response for certain compounds due to hyperactivation of transporters, causing cells in which Mad proteins are deleted to be significantly more resistant to several drugs.

This work establishes a novel chemical genomic means of interrogating transcriptional factors involved in PDR and applies this system to uncover multiple novel contributors to PDR regulation and a novel link between cell cycle regulation and PDR. These results not only increase our understanding of yeast biology but also provide novel targets for possible therapeutic intervention.

## Results

### Development of a biosensor system that couples PDR transporter expression to strain growth

One key challenge in dissecting the regulation of pleiotropic drug resistance in a high-throughput manner is linking the transcriptional response of PDR to a selectable phenotype, thereby allowing the system to be perturbed by genetic and chemical methods to determine the factors involved in PDR regulation. Toward this end, a plasmid-based biosensor that conferred a growth advantage to yeast cells upon exogenous chemical treatment and subsequent induction of transporter transcription was constructed (Figure 1A). This system consists of a yeast CEN/ARS plasmid on which the promoter of the PDR transporter being investigated (P*_PDR_*) was cloned upstream of the gene for imidazoleglycerol-phosphate dehydratase (*HIS3*), a protein essential for histidine biosynthesis, and the ***CYC1*** terminator (T*_CYC1_*). The promoters tested include those of *PDR5*, *SNQ2* and *YOR1*, three of the best-characterized ABC transporters in yeast. Strains transformed with biosensor constructs were grown in minimal defined media lacking histidine and containing varying concentrations of 3-Amino-1,2,4-triazole (3-AT, **4**), a potent and specific competitive inhibitor of His3 (16). The inclusion of an inhibitory but sublethal concentration of 3-AT in the growth media allows the system to be tuned for the different expression levels of each promoter being examined.

**Figure 1.**
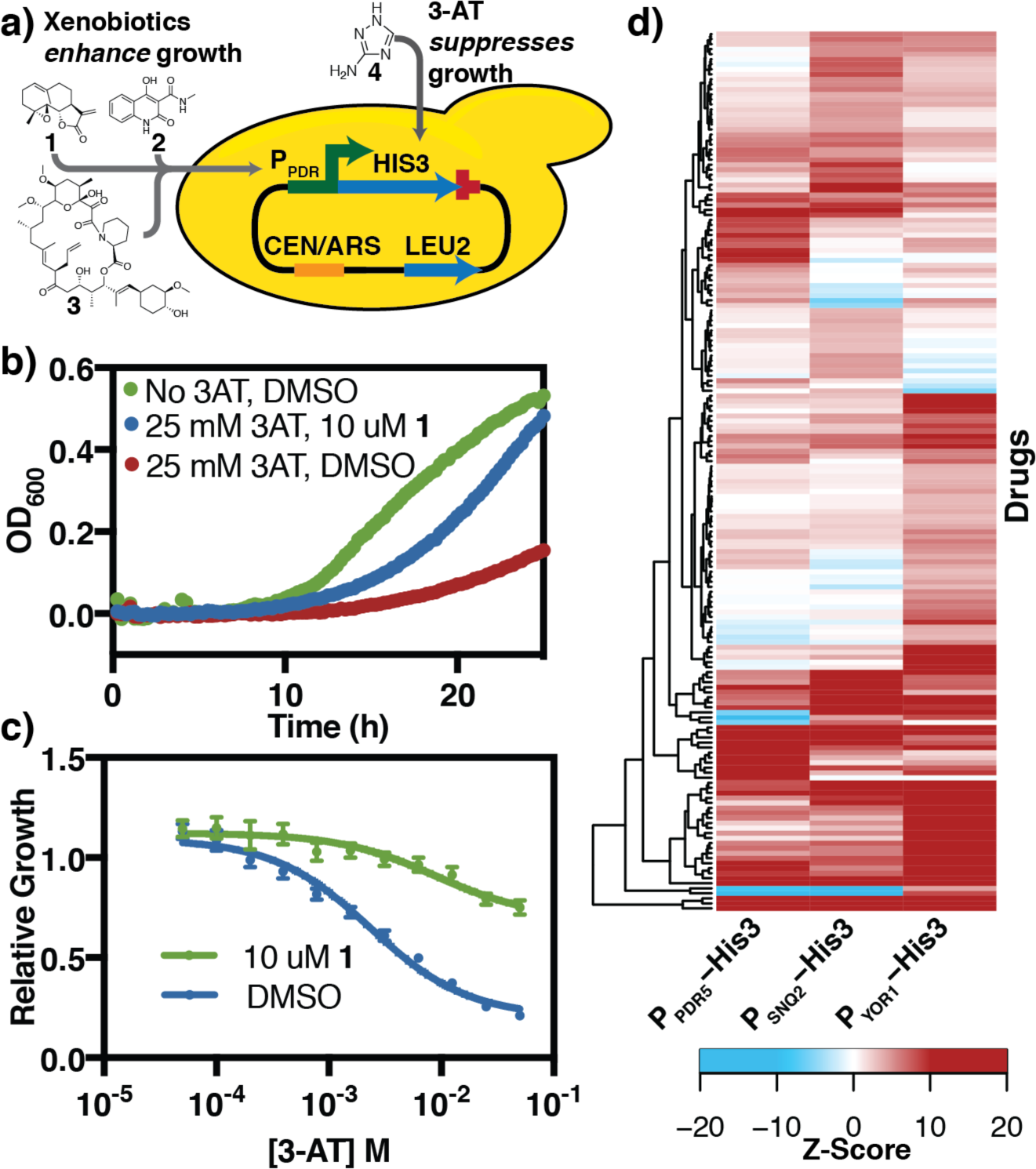
Construction of a biosensor system that couples PDR transporter expression and strain growth. (a) Schematic of the biosensor system and structures of inducing compounds used. The biosensor plasmids are outlined in Table S5. P*PDR* refers to promoters of PDR transporters, PDR5, SNQ2 or YOR1. Yeast strains transformed with these plasmids are grown in YNB media, without Leucine and Histidine. In addition, chemicals that induce the PDR response, such as Parthenolide (**1**), cbf_5236571 (**2**), and FK506 (**3**) were added to the media. Media also contained 3-AT(**4**), a potent competitive inhibitor of His3 to further control the growth and dynamic range of the system. (b) Growth curve of JLY31, a PDR5 biosensor strain showing growth suppression by **4** (25mM) and rescue by induction with **1** (10μM). Curves represent the average of 4 biological replicates. (c) Dose-response of JLY31 under increasing concentrations of **4**.Relative growth = AUC**4**/AUC_no_ **4**. Curves represent the best fit dose-response curve, and error bars represent SEM (n=4). (d) Heat map of drugs that significantly induces PDR transporters (Z≥3 for one or more biosensor construct). A survey of 800 natural products on the induction of JLY31-33 biosensor strain was conducted. A Z-score was calculated to reflect the strain growth, normalized to quality-control adjusted growth of the same strain treated with DMSO.

The utility of this system is demonstrated through by examining growth induction by a series of chemicals (Figure 1B, Figure S1A-C). In these experiments, a strain possessing the construct in which *HIS3* expression was driven by P_*PDR5*_ (JLY31) shows significantly increased growth in the presence of 3-AT upon treatment with 10 μM Parthenolide (**1**), a terpenoid natural product, as compared to DMSO control. This improved growth phenotype persisted in a dose-dependent manner across a range of 3-AT concentrations (Figure 1C). We observed similar growth induction upon treatment with **1** in strains carrying P_*SNQ2*_ (JLY32) and P_*YOR1*_ (JLY33) biosensor constructs (Figure S1D-K).

Previous studies demonstrated that the transcriptional induction of the PDR can vary significantly from compound to compound. Notably, FK506 (**3**), an immunosuppressant, induced *PDR5* and *SNQ2* expression, while cbf_5236571 (**2**) specifically induced *SNQ2* with little effect on *PDR5* (17). This biosensor system recapitulated these results, with **3** inducing growth in all three systems tested, while **2** strongly induces only the P_*SNQ2*_ and P_*YOR1*_ systems (Figure S1D-J) with growth being slightly suppressed in the P_*PDR5*_ case (Figure S1B, C).

To examine the generality of these results, we treated biosensor strains JLY31-33 with a library of 800 natural products (Figure 1D, Supplemental Data). For each strain-drug combination, a Z-score was calculated to reflect the change in strain growth upon treatment with drug, normalized to quality-control adjusted growth (see methods) of the same strain treated with DMSO. A high positive Z-score reflects a strong growth advantage under drug treatment and suggests the compound is a strong inducer of the specific PDR promoter. In contrast, a negative Z-score suggests a growth disadvantage, which can result from a combination of compound toxicity and little or no induction. We observed broad induction across varieties of compounds with 175 (22%) compounds leading to significant induction (Z>3) of at least one promoter. There was no apparent correlation between chemical structure and transporter induction for any of the three PDR promoters tested (Figure S2). The varied induction profiles observed for each pump in response to structurally diverse natural products further underscores the complexity of PDR regulation.

### Genome-wide, multiplexed interrogation of PDR reveals multiple candidate regulators of the PDR process

With a phenotypic screen for transporter induction established, we performed a genome-wide screens using the yeast deletion library to identify mutants that affect the transcriptional induction of PDR transporters. We transformed a barcoded yeast homozygous deletion collection with each of the P_*PDR5*_, P_*SNQ2*_, and P_*YOR1*_ biosensor constructs (pCH81-83). The pooled transformants were grown in media containing an inhibitory but sub-lethal concentration of 3-AT with either an inducing compound or DMSO as a control. The optimal concentrations of chemicals applied in each screen were determined through titrations that sampled biosensor response across a broad range of drug and 3-AT concentrations. After 6 generations of growth, the relative abundance of each deletion mutant in the treatment and control pools was quantified by microarray hybridization and analysis (Figure 2A). A strain overrepresented in the treatment group has a positive fold change (fc) and the gene deleted in this mutant is, therefore, a putative down-regulator of the induction of the promoter being examined. Conversely, negative fc values identified putative up-regulators of pump induction. A total of 9 screens were conducted: P_*PDR5*_, P_*SNQ2*_, and P_*YOR1*_ biosensors each with compounds **1**, **2**, and **3** (Figure 2B, Supplemental Data). These screens identified 314 hits, or deletions that were significantly different (log_2_(|fc|)>0.75 and a p<0.01) in at least one of the conditions tested.

**Figure 2.**
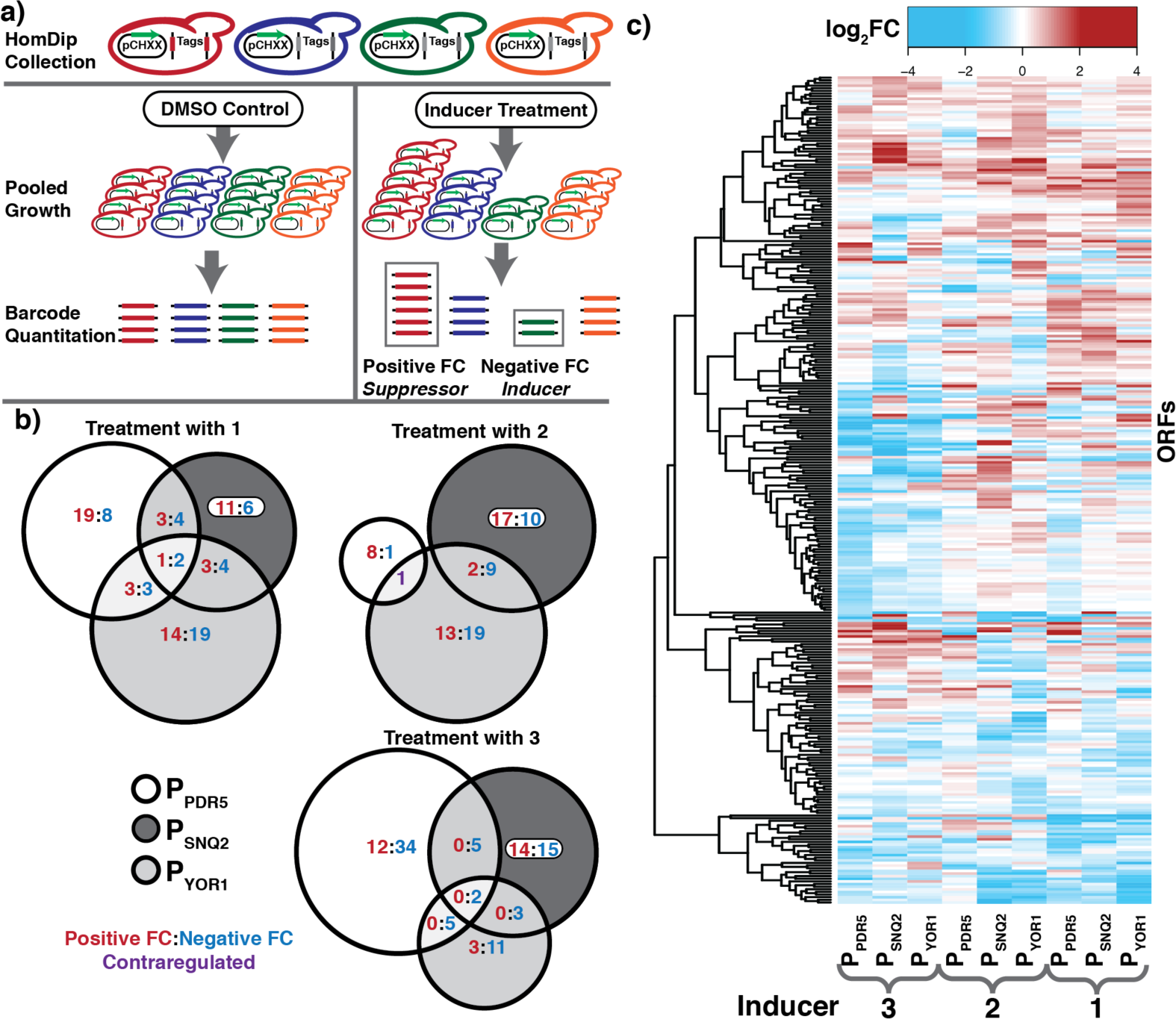
Multiplexed interrogation of PDR regulators reveals multiple candidate regulators of the PDR process. **(a)** Schematic of the homozygous deletion collection screening with biosensor system. Cultures were harvested after 6-generations of growth and relative abundance of each mutant in the treatment and control conditions were quantified by microarray analysis. **(b)** Venn Diagrams representing the overlapping hits across the 3 biosensor systems for each of the three inducers (**1-3**). Numbers represent ORFs that were significantly upregulated (red), downregulated (blue), or contraregulated (purple). **(c)** Heatmap of deletion mutants for which a log_2_(|fc|)>0.75 and p<0.01 for at least one condition. All data were calculated from 4 biological replicates.

While many deletion mutants only met the significance cutoff in a single condition, a number of hits appear general, demonstrating a modulation of multiple PDR proteins with multiple inducers (Figure 2C). We observed a relatively smaller number of hits in screening of P*_PDR5_* with **2**, and these hits have no overlap with P_SNQ2_ and P_YOR1_. This is to be expected as **2** does not induce a growth phenotype in the P*_PDR5_* system (Figure S1C). This suggests that this system has a low false positive rate and the false positives observed do not overlap with hits in other experiments.

Notably, the screens identified many known PDR regulators and factors that mediate cellular response to chemicals and stress. Yrr1, a known transcriptional regulator of Snq2 and Yor1, was identified as a hit in the screen with **2** on these two transporters (18). Analysis of gene ontology enrichment of hits identified in these screens shows enrichment for terms such as regulation of response to stimulus, regulation of response to stress and regulation of signaling, further demonstrating that the biosensor technology identified known elements of the PDR process (Table S1).

In addition to the known PDR-related pathways, several cellular processes were enriched in observed hits that were not previously associated with PDR, including mitotic SAC and cell cycle control, negative regulation of chromatin silencing, regulation of transcription, and regulation of primary metabolic process. To validate that the hits identified were truly modulating PDR induction and not artifacts of the biosensor system, we assayed the change in transcript levels upon xenobiotic treatment directly using quantitative PCR (qPCR). We focused the validation efforts on hits identified in two or more screens. Individual deletion mutants were treated for 1 hour with 50 µM of the compound to be screened during exponential growth. qPCR assays were performed to determine the fold induction of each PDR transporter (Figure 3, Table S2). Deletion of an up-regulator is expected to lead to less PDR induction, and thus a relative induction value lower than 1, while down-regulator deletion mutants are expected to have values higher than 1. The results of these individual experiments correlate well with those found in the growth assays, suggesting that the growth assays are a viable proxy for transcription (Table S2).

**Figure 3.**
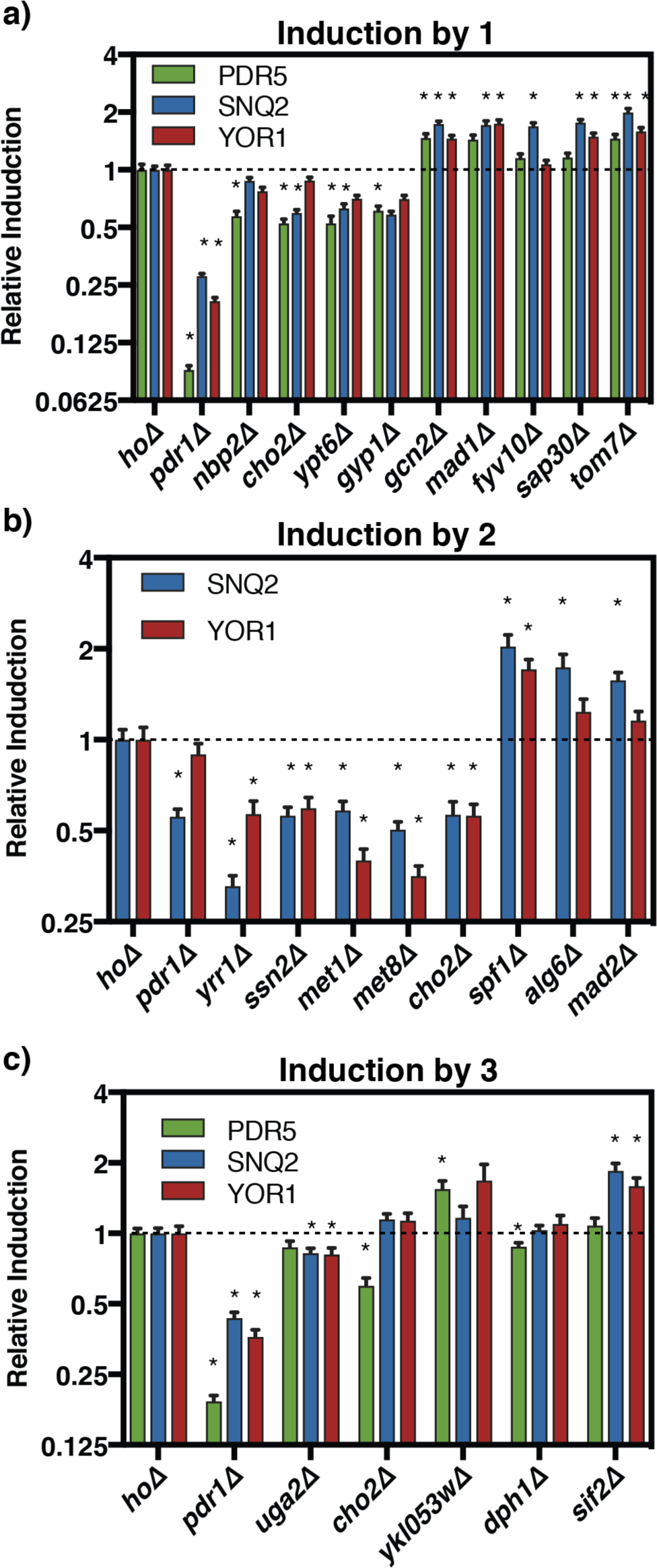
qPCR validation of PDR transcriptional responses confirms screen hits. Strains were grown to mid-exponential phase and treated with indicated compounds at 50 µM for 1 hr. Relative induction was calculated by comparing the fold induction of the transporter gene between *hoΔ* and the deletion strain. Error bars represent SEM (n=3). * indicates p<0.05, based on Student’s t-test on the fold induction of the transporter gene between *hoΔ* and the deletion strain. Treatment with (a)**1**, (b) **2**, and (c)**3**.

These validated hits have diverse cellular functions, with many not previously linked to the PDR. The qPCR assays confirmed that known transcriptional up-regulators, such as Pdr1 for all three transporters and Yrr1 for Snq2 and Yor1, are important for PDR transcription induction, with their deletion leading to significant decreases in pump induction. Similarly, we observed similar reductions with the deletion of factors involved in phospholipid metabolism (Cho2), cell wall integrity (Nbp2), and RNA polymerase II mediator complex (Ssn2), validating their important role in up-regulating PDR transcription. Additionally, select genes involved in methionine metabolism also appear to behave as transcriptional up-regulators of the PDR process (Met1, Met8).

For several up-regulators, these data correlate well with large-scale drug screens which demonstrated that deletion of these genes lead to increased drug sensitivity. For example, an *ssn2Δ* mutant was shown to have decreased resistance to drugs such as benzopyrene, chitosan, and geldanamycin and a *cho2Δ* mutant was suggested to have decreased resistance to drugs such as tellurite, benomyl and mycophenolic acid (19). Several membrane transporters appeared to act as down-regulators of PDR response, such as Tom7, a cellular transporter element involved in translocase of outer membrane complex, and Spf1, an ER membrane transporter important for intracellular membrane lipid composition. These qPCR results confirmed that the diverse groups of selected genes identified in the genome-wide screens are indeed regulators of the PDR process, and have established many new connections between distinct cellular process and PDR.

### Disruption of spindle assembly checkpoint leads to elevated PDR activation

In both growth-based screens and subsequent qPCR validations, we observed previously uncharacterized involvement of cell cycle regulators in the PDR transcriptional response. In particular, the components of mitotic SAC, Mad1-3, all appeared in growth screens and were validated by qPCR as down-regulators of the PDR process in response to exogenous drug treatment (Figures 2, 3). The mitotic SAC is a highly-conserved cell cycle surveillance mechanism that prevents abnormal chromosome segregation (20). While the exact mechanism of MAD proteins remains unknown, these proteins are generally believed to form complexes that inhibit anaphase promoting complex (APC/C), an E3 ubiquitin ligase that leads to the degradation of multiple downstream cell cycle proteins and enables sister-chromatid separation (21). Disruption of mitotic SAC leads to genome instability and has been implicated as the cause in many cancer cell lines (11) and inhibition of this complex has been proposed as a possible anticancer strategy. As deletion of genes for the MAD proteins leads to elevated PDR response, we set out to determine if *mad* deletion strains were more resistant to chemical treatment and, if so, whether PDR plays a role in conferring the resistance.

To identify compounds suitable for this assay, we mined previous chemogenomic datasets measuring fitness changes of the same deletion collection screened here in response to treatment with 3,250 small molecules (22). We identified a collection of structurally diverse compounds that induced less of a fitness defect when MAD genes were deleted as compared to DMSO control. These compounds include: cbf_5328528 (**5**), paf C16 (**6**), k035-0031 (**7**), 0180-0423 (**8**), N,N-dimethylsphingosine (**9**) (Figure 4A). We monitored the growth of reference *hoΔ* strain alongside the mad deletion strains, *mad1Δ*, *mad2Δ* and *mad3Δ* treatment with 140 µM **5**. The growth of all strains was significantly inhibited in this assay, but *mad* deletion strains grew better than *hoΔ* control (Figure 4B). The dose-response curves of mad deletion strains when treated with **5** are shifted significantly to the right as compared to *hoΔ*, and *mad* deletion strains have a statistically-significantly higher IC_50_ for **5**, demonstrating that mad deletion strains are significantly more resistant to this compound (Figure 4C, Table 1), consistent with the chemogenomic data. We observed a similar decrease in sensitivity of the *MAD* deletion mutants when treated with compounds **6**-**9**. Next, we performed qPCR analysis on these samples to determine if the transcription of PDR transporters was hyperactivated in the *MAD* deletion strains. **5** induces all three transporters and is a strong inducer of *PDR5* and *YOR1* (Table S3). We observed that all three transporters are significantly more activated in *MAD* deletion strains under **5** treatment (Figure 4F), suggesting a potential PDR involvement in the observed resistance. Similarly, compounds **6**-**9** show increased induction of at least one pump in at least one of the *MAD* deletion strains tested (Table S3).

**Figure 4.**
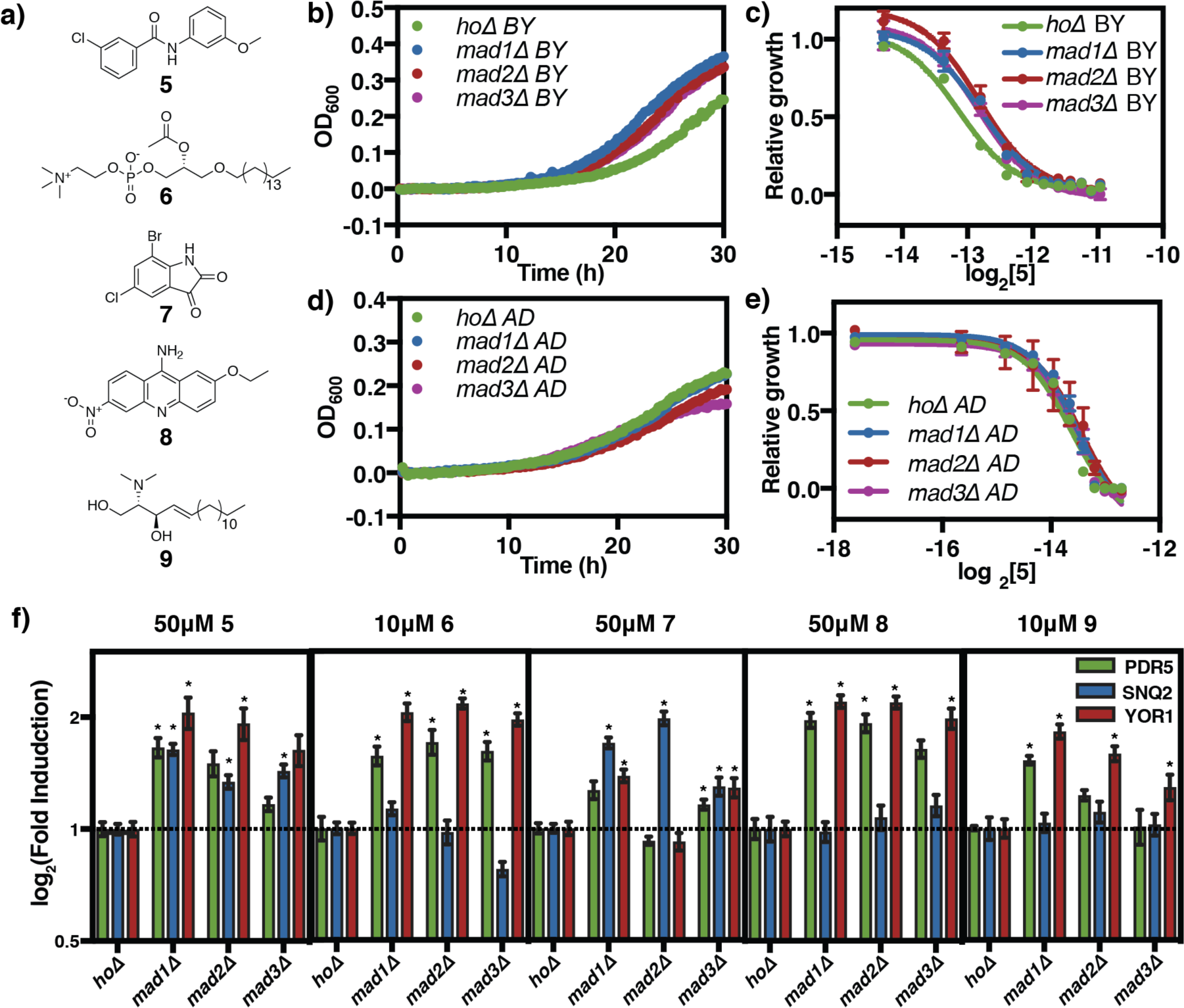
Disruption of spindle check point leads to elevated PDR activation. (a) Structures of compounds screened: cbf_5328528 (**5**), paf C-16 (**6**), k035-0031 (**7**), 0180-0423 (**8**), N,N-DMS (**9**). (b) Baseline corrected representative growth curve of *hoΔ* and *mad* deletion in BY4743 background under treatment with 140 µM **5**. OD_600_ were measured every 15 minutes over 30 hours. (c) Dose-response of strains in (b) over increasing **5** concentrations. Relative growth = AUC**5**/AUC_no drug_. All wells contain the same DMSO concentration. AUC calculation was performed with baseline corrected growth curve. Line represents best fit dose-response curve, and error bar represents SEM (n=3). (d) Baseline corrected representative growth curve of *hoΔ* and *mad* deletion in AD1-9 PDR transporter null background under 77.5 µM **5** treatment. (e) Dose-response of strains in (d) over increasing **5** concentrations. (f) qPCR validation of transcription induction of all three transporters under the treatment of all five compounds in (a). Methods are identical to those in Figure 3. Error bars represent SEM (n=3). * indicates p<0.05.

**Table 1:** IC_50_ of mad deletions strains in BY4743 and AD1-9 background. IC_50_ were calculated based on best-fitting curve displayed. P-values were calculated using a student’s T-test with 3 biological replicates. Significant p-values (<0.05) are italicized in bold.

| Drug | Strain | BY4743 |  | AD1-9 |  |
| --- | --- | --- | --- | --- | --- |
| | | IC<sub>50</sub>(μM) | p | IC<sub>50</sub>(μM) | p |
| 5 | *ho*Δ | 111.6 | - | 78.3 | - |
| | *mad1*Δ | 140.2 | ***0.009*** | 86.3 | 0.216 |
| | *mad2*Δ | 138.3 | ***0.015*** | 90.0 | 0.286 |
| | *mad3*Δ | 137.3 | ***0.018*** | 88.1 | 0.180 |
| 6 | *ho*Δ | 8.6 | - | 7.84 | - |
| | *mad1*Δ | 11.7 | ***0.018*** | 7.32 | 0.520 |
| | *mad2*Δ | 12.4 | ***0.014*** | 7.02 | 0.327 |
| | *mad3*Δ | 11.6 | ***0.021*** | 5.86 | 0.144 |
| 7 | *ho*Δ | 53.5 | - | 43.76 | - |
| | *mad1*Δ | 80.5 | ***0.013*** | 41.11 | 0.637 |
| | *mad2*Δ | 73.6 | ***0.031*** | 45.94 | 0.737 |
| | *mad3*Δ | 80.0 | ***0.013*** | 39.71 | 0.426 |
| 8 | *ho*Δ | 252.8 | - | 0.95 | - |
| | *mad1*Δ | 359.9 | ***0.007*** | 0.59 | 1.000 |
| | *mad2*Δ | 345.3 | ***0.020*** | 0.59 | 0.470 |
| | *mad3*Δ | 326.6 | ***0.014*** | 0.63 | 0.917 |
| 9 | *ho*Δ | 12.3 | - | 0.60 | - |
| | *mad1*Δ | 16.5 | ***0.021*** | 8.64 | 0.909 |
| | *mad2*Δ | 16.4 | ***0.019*** | 7.90 | 0.533 |
| | *mad3*Δ | 19.9 | ***0.005*** | 7.17 | 0.176 |

To ascertain if the increased expression of the PDR transporters is responsible for the increased resistance, we constructed the same *hoΔ*, *mad1Δ*, *mad2Δ*, and *mad3Δ* deletions in the AD1-9 background (named *hoΔ* AD, *mad1Δ*, *2Δ*, *3Δ* AD, respectively). AD1-9 is a strain in which the genes for 9 plasma-membrane bound PDR transporters, including *PDR5*, *SNQ2* and *YOR1*, are knocked out (23). If the observed increased resistance is due to increased expression of PDR transporters, we expected that the increased resistance phenotype will not persist in this background.

This hypothesis was confirmed in repeating the growth inhibition experiment in AD1-9 background. In these strains, both *ho* and *mad* deletions strains are again inhibited by **5**, but the *mad* deletion strains with transporters knocked out no longer exhibit the increased resistance (Figure 4D). The **5** dose-response curves of mad deletion strains are not significantly different from that of *hoΔ* AD, and the IC_50_s of these strains are also not significantly different, suggesting these strains have a similar sensitivities to **5** (Figure 4E, Table 1). It is also notable that AD1-9 strains are more sensitive to **5**, suggesting a PDR involvement in the resistance to this compound (Figure 4F). To ensure this observation is not specific to a particular compound, we performed the same set of experiments with compounds **6**-**9**. In all cases, deletion of *MAD* genes strains led to elevated resistance and increased PDR transporter induction. This increased resistance is no longer observed when PDR transporters are knocked out in the AD1-9 (Figure 4F, Table 1, Figure S3, Table S3). These results demonstrate that disruption of mitotic SAC complex leads to elevated PDR activation and increased drug resistance, revealing a novel link between cell cycle regulation and pleiotropic drug resistance.

## Discussion

In this study, we developed a biosensor technology that couples transcriptional induction of yeast pleiotropic drug response to growth rate, an approach that builds upon previous approaches with fluorescence-based reporters (24) by allowing screening of mutant pools in competitive growth assays. This system specifically captured dose-dependent PDR transcriptional response induced by a variety of compounds. Pooled screens of the homozygous diploid yeast deletion collection identified 314 putative PDR regulators spanning a broad range of functional areas, including, but not limited to, response to chemical stimulus, lipid metabolism, translation, RNA metabolism, and cell cycle. We subsequently confirmed 20 genes that had been identified in multiple screens as transcriptional regulators of the PDR process by demonstrating that their deletion directly affects transcript abundance upon exogenous chemical treatment. In particular, we discovered mitotic spindle checkpoint factors, Mad1-3, as down-regulators of the PDR process. Deletion of these genes leads to hyperactivation of PDR transporters upon exogenous compound treatment, leading to elevated resistance.

The involvement of cell cycle regulators in PDR response has not been previously established, but is consistent with several previous observations. It was initially proposed more than a hundred years ago that defects in proper chromosomal segregation are tumorigenic (25). Subsequent work suggested that mutations in the SAC mechanism are important reasons for aneuploidy and cancer (20). Many of these cells with impaired SAC function showed increased resistance to mitotic inhibitors in survival or growth assays, both in artificially induced conditions or naturally occurring cancer cell lines (26, 27). For instance, in a study with multidrug-resistant colon cancer cell lines where the expression of Mad2 was suppressed by more than 50%, a significantly enhanced PDR expression level was observed, leading the authors to suggest that elevated PDR in these SAC impaired cells is the reason for increased drug resistance (28).

This study builds upon these observations, demonstrating that PDR activation due to SAC impairment could be another mechanism that cancer cells, many of which have mutations of SACs, becomes resistant to drug treatment. Mitotic proteins have been long pursued as cancer drug targets (29). More recent development efforts have focused on kinesins such as CENP-E, KSP and kinases such as Plk1, Aurora A and B, Mps1. Among these new targets, targeting Aurora B and Mps1 are directly targeting SAC inhibition (29). These results suggest additional considerations when thinking of the SAC members as drug targets. Specifically, by restoring the function of Mad1-3, cancers cells may not only have better regulated mitosis and proliferation, but may also become less resistant to traditional chemotherapies.

In addition to those involved in cell cycle control, this work identified multiple other previously uncharacterized PDR regulators, many of which participate in processes that have a known connection to the PDR. For example, we have noticed SPF1 and TOM7, two genes related to transportation channels on mitochondrial outer membrane, as PDR regulators (30, 31). Defects in these factors lead to mitochondrial damage, which can activate retrograde signaling. In a PDR3-dependent manner, retrograde signaling has been shown to then induce transcriptional activation of PDR genes to facilitate transporting damaging species outside the cells (32).

Another factor that lies in the intersection of PDR and other pathways is Cho2, encoding the phosphatidylethanolamine methyltranferase in phospholipids synthesis, which is involved in the cell wall integrity (CWI) pathway (33). In response to certain chemical stress, such as organic solvents, PDR and CWI were activated in a coordinated fashion to cope with the stress, suggesting potential co-regulation and crosstalk between these stress response pathways (34). Identification of Ssn2, a subunit of RNA polymerase II mediator complex, and Gcn2, a kinase for α-subunit of eIF2, as regulators of the PDR process is not unexpected as transcriptional regulation can have significant impact on gene expression (35). In both human cell lines and *Candida albicans*, chromatin-remodeling complex, which Ssn2 and Gcn2 interact with, was observed to control expression of PDR transporters (36, 37).

In this study, we have presented the genome-wide dissection of the yeast PDR using a series of biosensors responsive to a variety of diverse small molecules. Through this work, we have confirmed the role of several proteins in PDR regulation and identified multiple additional factors with significant but previously uncharacterized regulatory roles. Further, we have demonstrated that, in the presence of several toxic compounds, disruption of mitotic spindle checkpoint assembly leads to elevated PDR and increased resistance. These results not only demonstrate the biosensor system as a viable tool to investigate PDR, but also uncover novel control of the process and a connection to cell cycle regulation.

## Materials and Methods

### Media and growth conditions

Two *S. cerevisiae* strains were primarily used in this study: BY4743 (*MAT*a/α *his3*Δ*1/his3*Δ*1 leu2*Δ*0/leu2*Δ*0 LYS2/lys2*Δ*0 met15*Δ*0/MET15 ura3*Δ*0/ura3*Δ*0*) and AD1-9 (MATα, pdr1-3, ura3, his1, ∆yor1::hisG, ∆snq2::hisG, pdr5-∆2::hisG, ∆pdr10::hisG, ∆pdr11::hisG, ∆ycf1::hisG, pdr3-∆2::hisG, ∆pdr15::hisG, pdr1-∆3::hisG) (23). Yeast was grown in YNB (yeast nitrogen base) media in all experiments. YNB media contains 1.7 g/L yeast nitrogen base (MP Biomedicals, catalog no. 114027512), 5 g/L ammonium sulfate, with selected amino acid based on specific experiments in the following concentrations: 20 mg/L histidine, 60 mg/L leucine, and 20 mg/L uracil. This media was further supplemented with 3-Amino-1,2,4-triazole (3-AT) at various concentrations as indicated experiments.

### Chemical reagents

The Microsource Pure Natural Products library (MSNP) was purchased from Discovery Systems. Parthenolide, 3-AT, paf C-16 and N,N-DMS were purchased from Sigma-Aldrich (catalog no. P0667, 61-82-5, P4904, and SML0311, respectively). FK506 was purchased from Enzo Life Sciences (catalog no. ALX-380-008). Cbf_5236571 and cbf_5328528 were purchased from ChemBridge. K035-0031 was purchased from Enamine (catalog no. 50-138-184). 0180-0423 was purchased from ChemDiv.

### Growth rate analysis of isogenic cultures and generation of dose response curve

All growth experiments were conducted with GENios microplate reader system (Tecan) in which optical density (OD_600_) was measured every 15 minutes over the entire course of the experiment. For experiments with isogenic strains, all growth assays were conducted in Nunc MicroWel 96-Well Microplates (Thermo Scientific) with 100 uL in each well. Pooled screens were conducted in Corning Costar 48-well cell culture plate (Sigma-Aldrich). Strains were inoculated into YNB media overnight, diluted into fresh media to allow for additional growth for 4 – 6 hours, when they are in exponential growth, and then diluted to OD_600_=0.01 to start the growth assays. Indicated concentration of various drugs were added to plates using a Tecan D300e Digital Dispenser.

Relative growth was determined as previously reported (17). Briefly, OD_600_ was measured every 15 minutes during the experiment. The mean of the first 10 measurements was used to baseline correct all subsequent measurements. Unless otherwise stated, all AUC measurements reported here are the sum of the baseline corrected OD_600_s over a 30 hour period.

For dose-response curves, relative growth values for each condition were computed to compare the rate of growth of the same strain under the drug treatment vs. no drug/DMSO control: Relative growth = AUC_drug_/AUC_DMSO_. Modelling was completed in GraphPad Prism version 7.0 for Windows, GraphPad Software, La Jolla California USA, following standard practice.

### Screen of MSNP collection

BY4743 *hoΔ*, was transformed with each of pCH81, pCH82, and pCH82 (Table S5). Strains were grown to stationary phase overnight in minimal media overnight and diluted to OD_600_ = 0.01 into minimal media containing an inhibitory concentration of 3-AT (100 mM for strains with pCH81, 50 mM for strains with pCH82 and 2 mM for strains containing pCH83). 100 uL of diluted culture was placed into each well of a 96 well plate well. 1 ul of each compound from all 800 compounds from the MSNP collection was inoculated into the growth media, leading to 1:100 dilution from the stock concentration. OD_600_ was monitored every 15 minutes for 30 hours. Each 96-well plate included 16 DMSO controls. Compound induction was quantified by calculating the following Z-score: Z = [AUC_drug_-AUC_mean(DMSO)_]/AUC_SD(DMSO)_. Compounds with a Z-score >3 were deemed to be significant inducers.

### Screening of yeast homozygous deletion collection

To construct a homozygous deletion collection that contains the biosensor constructs, pCH81-83 were transformed into the pooled *S. cerevisiae* deletion collection using a standard lithium acetate protocol (38, 39). Each transformation resulted approximately 10^5^ transformants that were subsequently pooled together. Pooled transformants were grown overnight to saturation and diluted to OD_600_ = 0.01. Diluted cultures were left to recover for 4 hours prior to challenge with both inducer and 3-AT which were then added in the following concentrations: 200 µM 3-AT with 10 µM **1** for all three promoters, 200 µM 3-AT and 10 µM **2** for all three promoters, 800 µM 3-AT and 10 µM **3** for P*_PDR5_*, 1.6 mM 3-AT and 10 µM **3** for P*_SNQ2_*, and 320 µM 3-AT and 10 µM **3** for P*_YOR1_*. 700 μl of culture was grown in each well of a 48 well plate at 30 °C with orbital shaking in Infinite plate readers (Tecan).

Growth of the pooled culture was monitored every 15 minutes. To maintain cultures in log phase throughout the growth experiment, after 3 generations of competitive growth, cultures were diluted to OD_600_= 0.075, and grown for a further 3 generations after which 600 μl of culture was harvested saved to a 4 °C cooling station (Torrey Pines). This amounted to approximately 6 culture doublings, or 6 generations of growth, from the beginning of the experiment. Pipetting events were triggered automatically by Pegasus Software and performed by a Freedom EVO workstation (Tecan).

Genomic DNA from the pools was prepared using a YeastStar genomic DNA prep (Zymo Research) and deletion collection barcodes were amplified as previously reported (16). Barcode quantitation was performed using Genflex Tag 16K Array v2 chip (Affymetrix) by following standard procedures (38, 39). Relative abundance of each barcode in induced vs. control conditions treated with equivalent amount of DMSO was used to determine induction of the promoter being examined, as previously described (17). All p-values were determined by a Student’s t-test for 4 biological replicates.

### Quantitative RT-qPCR experiments

In experiments examining the transcriptional response of PDR promoters, individual strains were grown to saturation in minimal media overnight. Cultures were then diluted to OD_600_=0.2 in fresh minimal media and grown for a further 4 hours. Cells were then treated with drugs at the inducing concentration (Figures 3, 4F) for 1 hour. 2 OD-equivalents of cells were then harvested for whole cell RNA extraction and DNA removal using RiboPure RNA Purification Kit (ThermoFisher, catalog no. AM1926). Purified RNA samples were reverse transcribed to single-stranded cDNA using High-Capacity RNA-to-cDNA Kit (ThermoFisher, catalog no. 4387406). These cDNA samples were used as template for quantitative PCR experiments. These q-PCR reactions included SYBR Green PCR Master Mix (ThermoFisher, catalog no. 4309155), and primers at 5 µM concentration. Primers were designed with the primer3 software package and were tested in previously published work (17). They are expected to yield ~120 bp amplicons for PDR promoters and ACT1 (control). qPCR reactions were performed on 7900HT Fast Real-Time PCR System (ThermoFisher) in 384-well format. qPCR data were analyzed using software integrated to the 7900HT system using automatic threshold determination. ΔCt values for each transport gene were calculated for each set of samples by comparing the difference between the Ct value of sample same treated with drug vs. DMSO control: ΔCt_transporter_=Ct_drug_-Ct_DMSO_. This ΔCt_transporter_ value is then normalized for cell numbers by comparing it with ΔCt_ACT1_: ΔΔCt_transporter_ = ΔCt_ACT1_ – ΔCt_transporter_. Student t-tests were applied on ΔΔCt level by comparing the magnitude of ΔΔCt in a deletion strain to that in *hoΔ* control. In Figures 3 and 4, a ΔΔΔCt_transporter_ was further calculated by computing the difference between ΔΔCt_transporter_ in a specific deletion strain and that in the *hoΔ* strain: ΔΔΔCt_transporter, deletion strain_ = ΔΔCt_transporter, deletion strain_ - ΔΔCt_transporter, *hoΔ*_.

### Construction of AD1-9 deletion strains

AD1-9 deletion background strain was a previous creation with all transporters individually knocked out (23). Further, HO, MAD1-3 genes are knocked out in this background using genetic constructs and methods that were used to create the yeast deletion collection (40). Briefly, primer pairs that amplify regions that encode *Δmad1-3::KanMX4* or on *Δho::KanMX4* respective BY4743 deletion strains were used to amplify knockout cassettes, with homologous regions on both end of the amplicon. This amplicon was then transformed into AD1-9 background strain, and the transformants were selected for kanamycin resistance, followed by sequencing of targeted region to confirm integration identity. The resulted stains contain *Δmad1-3::KanMX4* or on *Δho::KanMX4* on AD1-9 background.

## Acknowledgement

We thank Angela Chu for thoughtful discussion and help in deletion collection. This work was supported by NIH grant U01 GM110706.

## Data Availabiltiy

All microarray data from genome-wide screens is available from EMBL-EBI ArrayExpress under accession number E-MTAB-6655.

